# Attentional Modulation of Hierarchical Speech Representations in a Multitalker Environment

**DOI:** 10.1101/2020.12.05.412957

**Authors:** Ibrahim Kiremitçi, Özgür Yilmaz, Emin Çelik, Mo Shahdloo, Alexander G Huth, Tolga Çukur

## Abstract

Humans are remarkably adept in listening to a desired speaker in a crowded environment, while filtering out non-target speakers in the background. Attention is key to solving this difficult cocktail-party task, yet a detailed characterization of attentional effects on speech representations is lacking. It remains unclear across what levels of speech features and how much attentional modulation occurs in each brain area during the cocktail-party task. To address these questions, we recorded whole-brain BOLD responses while subjects either passively listened to single-speaker stories, or selectively attended to a male or a female speaker in temporally-overlaid stories in separate experiments. Spectral, articulatory, and semantic models of the natural stories were constructed. Intrinsic selectivity profiles were identified via voxelwise models fit to passive listening responses. Attentional modulations were then quantified based on model predictions for attended and unattended stories in the cocktail-party task. We find that attention causes broad modulations at multiple levels of speech representations while growing stronger towards later stages of processing, and that unattended speech is represented up to the semantic level in parabelt auditory cortex. These results provide insights on attentional mechanisms that underlie the ability to selectively listen to a desired speaker in noisy multi-speaker environments.

## INTRODUCTION

Humans are highly adept at perceiving a target speaker in crowded multi-speaker environments (Shinn-Cunningham and Best 2008; Kidd and Colburn 2017; Li et al. 2018). Auditory attention is key to behavioral performance in this difficult “cocktail-party problem” (Cherry 1953; Fritz et al. 2007; McDermott 2009; Bronkhorst 2015; Shinn-Cunningham et al. 2017). Literature consistently reports that attention selectively enhances cortical responses to the target stream in auditory cortex and beyond, while filtering out non-target background streams (Hink and Hillyard 1976; Teder et al. 1993; Alho et al. 1999, 2003, 2014; Jäncke et al. 2001, 2003; Lipschutz et al. 2002; Rienne et al. 2008, 2010; Elhilali et al. 2009; Gutschalk and Dykstra 2014). However, the precise link between the response modulations and underlying speech representations is less clear. Speech representations are hierarchically organized across multiple stages of processing in cortex, with each stage selective for diverse information ranging from low-level acoustic to high-level semantic features (Davis and Johnsrude 2003; Griffiths and Warren 2004; Hickok and Poeppel 2004, 2007; Rauschecker and Scott 2009; Okada et al. 2010; Friederici 2011; Di Liberto et al. 2015; de Heer et al. 2017; Brodbeck et al. 2018a). Thus, a principal question is to what extent attention modulates these multi-level speech representations in the human brain during a cocktail-party task (Miller 2016; Simon 2017).

Recent electrophysiology studies on the cocktail-party problem have investigated attentional response modulations for natural speech stimuli (Kerlin et al. 2010; Ding and Simon 2012a, 2012b; Mesgarani and Chang 2012; Power et al. 2012; Zion Golumbic et al. 2013; Puvvada and Simon 2017; Brodbeck et al. 2018b; O’Sullivan et al. 2019; Puschman et al. 2019). Ding and Simon (2012a, 2012b) fit spectrotemporal encoding models to predict cortical responses from the speech spectrogram. Attentional modulation in the peak amplitude of spectrotemporal response functions was reported in planum temporale in favor of the attended speech. Mesgarani and Chang (2012) built decoding models to estimate the speech spectrogram from responses measured during passive listening and examined the similarity of the decoded spectrogram during a cocktail-party task to the isolated spectrograms of attended versus unattended speech. They found higher similarity to attended speech in non-primary auditory cortex. Zion Golumbic et al. (2013) reported amplitude modulations in speech-envelope response functions towards attended speech across auditory, inferior temporal, frontal and parietal cortices. Other studies using decoding models have similarly reported higher decoding performance for the speech envelope of the attended stream in auditory, prefrontal, motor and somatosensory cortices (Puvvada and Simon 2017; Puschmann et al. 2019). Brodbeck et al. (2018b) further identified peak amplitude response modulations for sub-lexical features including word onset and cohort entropy in temporal cortex. Note that because these electrophysiology studies fit models for acoustic or sub-lexical features, the reported attentional modulations primarily comprised relatively low-level speech representations.

Several neuroimaging studies have also examined whole-brain cortical responses to natural speech in a cocktail-party setting (Nakai et al. 2005; Alho et al. 2006; Ikeda et al. 2010; Hill and Miller 2010; Wild et al. 2012; Regev et al. 2019; Wikman et al. 2021). In the study of Hill and Miller (2010), subjects were given an attention cue (attend to pitch, attend to location or rest) and later exposed to multiple speech stimuli where they performed the cued task. Partly overlapping frontal and parietal activations were reported, during both the cue and the stimulus exposure periods, as an effect of attention to pitch or location in contrast to rest. Furthermore, pitch-based attention was found to elicit higher responses in bilateral posterior and right middle superior temporal sulcus, whereas location-based attention elicited higher responses in left intraparietal sulcus. In alignment with electrophysiology studies, these results suggest that attention modulates relatively low-level speech representations comprising paralinguistic features. In a more recent study, Regev et al. (2019) measured responses under two distinct conditions: while subjects were presented bimodal speech-text stories and asked to attend to either the auditory or visual stimulus, and while subjects were presented unimodal speech or text stories. Inter-subject response correlations were measured between unimodal and bimodal conditions. Broad attentional modulations in response correlation were reported from primary auditory cortex to temporal, parietal and frontal regions in favor of the attended modality. While this finding raises the possibility that attention might also affect representations in higher-order regions, a systematic characterization of individual speech features that drive attentional modulations across cortex is lacking.

An equally important question regarding the cocktail-party problem is whether unattended speech streams are represented in cortex despite the reported modulations in favor of the target stream (Bronkhorst 2015; Miller 2016). Electrophysiology studies on this issue identified representations of low-level spectrogram and speech envelope features of unattended speech in early auditory areas (Ding and Simon 2012a, 2012b; Mesgarani and Chang 2012; Zion Golumbic et al. 2013; Puvvada and Simon 2017; Brodbeck et al. 2018b; Puschmann et al. 2019), but no representations of linguistic features (Brodbeck et al. 2018b). Meanwhile, a group of neuroimaging studies found broader cortical responses to unattended speech in superior temporal cortex (Scott et al. 2004, 2009a; Wild et al. 2012; Scott and McGettigan 2013; Evans et al. 2016; Regev et al. 2019). Specifically, Wild et al. (2012) and Evans et al. (2016) reported enhanced activity associated with the intelligibility of unattended stream in parts of superior temporal cortex extending to superior temporal sulcus. Although this implies that responses in relatively higher auditory areas carry some information regarding unattended speech stimuli, the specific features of unattended speech that are represented across the cortical hierarchy of speech is lacking.

Here we investigated whether and how attention affects representations of attended and unattended natural speech across cortex. To address these questions, we systematically examined multi-level speech representations during a diotic cocktail-party task using naturalistic stimuli. Whole-brain BOLD responses were recorded in two separate experiments (Figure 1) while subjects were presented engaging spoken narratives from *The Moth Radio Hour.* In the passive-listening experiment, subjects listened to single-speaker stories for over two hours. Separate voxelwise models were fit that measured selectivity for spectral, articulatory, and semantic features of natural speech during passive listening (de Heer et al. 2017). In the cocktail-party experiment, subjects listened to temporally-overlaid speech streams from two speakers while attending to a target category (male or female speaker). To assess attentional modulation in functional selectivity, voxelwise models fit during passive listening were used to predict responses for the cocktail-party experiment. Model performances were calculated separately for attended and unattended stories. Attentional modulation was taken as the difference between these two performance measurements. Comprehensive analyses were conducted to examine the intrinsic complexity and attentional modulation of multi-level speech representations and to investigate up to what level of speech features unattended speech is represented across cortex.

**Figure 1:**
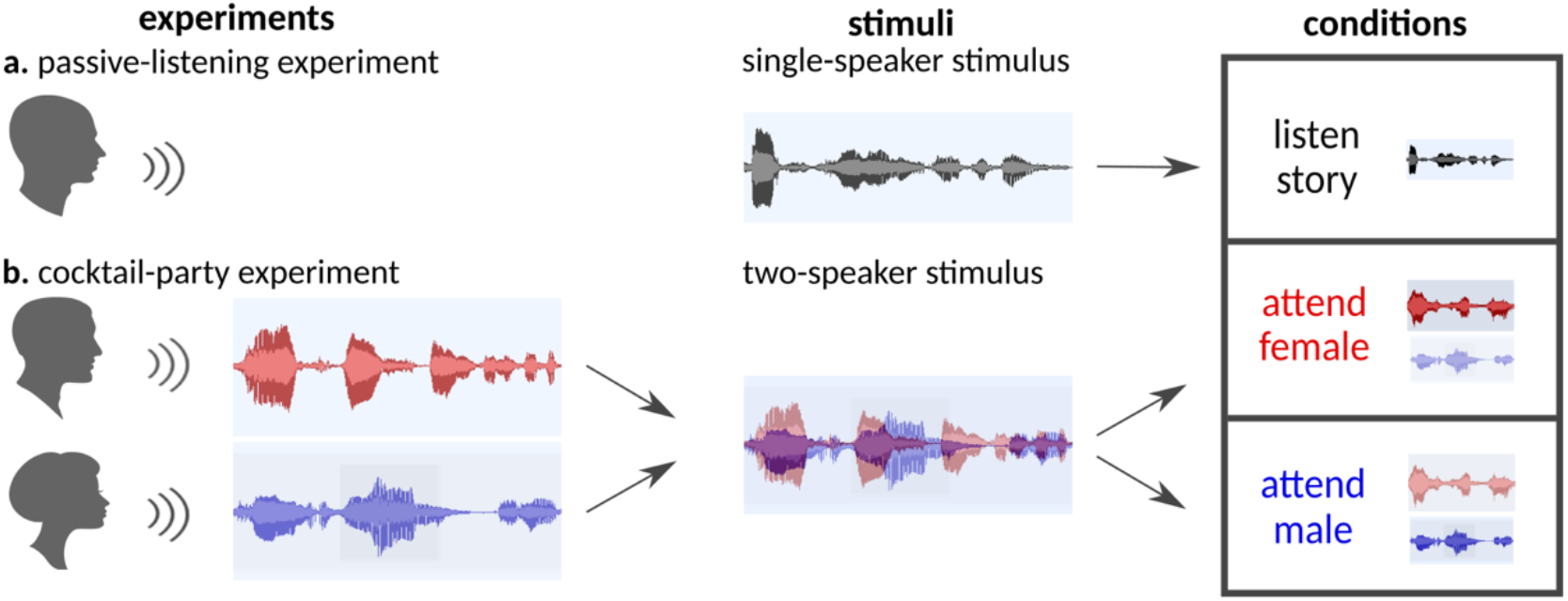
Experimental design. **a.** *Passive-listening experiment.* 10 stories from Moth-Radio-Hour were used to compile a single-speaker stimulus set. Subjects were instructed to listen to the stimulus vigilantly without any explicit task in the passive-listening experiment. **b.** *Cocktail-party experiment.* A pair of stories told by individuals of different genders were selected from the single-speaker stimulus set and overlaid temporally to generate a two-speaker stimulus set. Subjects were instructed to attend either to the male or female speaker in the cocktail-party experiment. The same two-speaker story was presented twice in separate runs while the target speaker was varied. Attention condition was fixed within runs and it alternated across runs.

## Materials and Methods

### Participants

Functional data were collected from five healthy adult native subjects (four males and one female; aged between 26 and 31) who had no reported hearing problems and were native English speakers. The experimental procedures were approved by the Committee for the Protection of Human Subjects at University of California, Berkeley. Written informed consent was obtained from all subjects.

### Stimuli

Figure 1 illustrates the two main types of stimuli used in the experiments: single-speaker stories and two-speaker stories. Ten single-speaker stories were taken from The Moth Radio Program: “Alternate Ithaca Tom” by Tom Weiser; “How to Draw a Nekkid Man” by Tricia Rose Burt; “Life Flight” by Kimberly Reed; “My Avatar and Me” by Laura Albert; “My First Day at the Yankees” by Matthew McGough; “My Unhurried Legacy” by Kyp Malone; “Naked” by Catherine Burns; “Ode to Stepfather” by Ethan Hawke; “Targeted” by Jen Lee and “Under the Influence” by Jeffery Rudell. All stories were told before a live audience by a male or female speaker, and they were about 10-15 min long. Each two-speaker story was generated by temporally overlaying a pair of stories told by different genders and selected from the single-speaker story set. When the durations of the two single-speaker stories differed, the longer story was clipped from the end to match durations. Three two-speaker stories were prepared: from “Targeted” and “Ode to Stepfather” (cocktail1); from “How to Draw a Nekkid Man” and “My First Day at the Yankees” (cocktail2); and from “Life Flight” and “Under the Influence” (cocktail3). In the end, the stimuli consisted of ten single-speaker and three two-speaker stories.

### Experimental procedures

Figure 1 outlines the two main experiments conducted in separate sessions: passive-listening and cocktail-party experiments. In the passive-listening experiment, subjects were instructed to listen to single-speaker stories vigilantly without an explicit attentional target. To facilitate sustained vigilance, we picked engaging spoken narratives from the Moth Radio Hour (see Stimuli). Each of the ten single-speaker stories was presented once in a separate run of the experiment. Two two-hour sessions were conducted, resulting in ten runs of passive-listening data for each subject. In the cocktail-party experiment, subjects were instructed to listen to two-speaker stories while attending to a target speaker (either the male or the female speaker). Our experimental design focuses on attentional modulations of speech representations when a stimulus is attended versus unattended. Each of the three cocktail-stories was presented twice in separate runs. This allowed us to present the same stimulus set in attended and unattended conditions to prevent potential biases due to across-condition stimulation differences. To minimize adaptation effects, different two-speaker stories were presented in consecutive runs while maximizing the time window between repeated presentations of a two-speaker story. Attention condition alternated across consecutive runs. An exemplary sequence of runs was: cocktail1-M (attend to male speaker in cocktail1), cocktail2-F (attend to female speaker in cocktail2), cocktail3-M, cocktail1-F, cocktail2-M, and cocktail3-F. The first attention condition assigned to each two-speaker story (M or F) was counterbalanced across subjects. This resulted in a balanced assignment of ‘attended’ versus ‘unattended’ conditions during the second exposure to each two-speaker story. Furthermore, for each subject, the second exposure to half of the single-speaker stories (3 out of 6 included within the two-speaker stories) coincided with the ‘attended’ condition, whereas the second exposure to the other half coincided with the ‘unattended’ condition. Hence, second exposure to each story was balanced across ‘attended’ and ‘unattended’ conditions both within and across subjects. A two-hour session was conducted, resulting in six runs of cocktail-data. Note that the two-speaker stories used in the cocktail-party experiment were constructed from the single-speaker story set used in passive-listening experiment. Hence, for each subject, the cocktail-party experiment was conducted several months (∼5.5 months) after the completion of the passive-listening experiment to minimize potential repetition effects. The dataset collected from the passive-listening experiment was previously analyzed (Huth et al. 2016; de Heer et al. 2017); however, the dataset collected from the cocktail-party experiment was specifically collected for this study.

In both experiments, the length of each run was tailored to the length of the story stimulus with additional 10 sec of silence both before and after the stimulus. All stimuli were played at 44.1 kHz and delivered binaurally to both ears using Sensimetrics S14 in-ear piezo-electric headphones. The Sensimetrics S14 is an MRI-compatible auditory stimulation system with foam canal tips to reduce scanner noise (above 29 dB as stated in specifications). The frequency response of the headphones was flattened using a Behringer Ultra-Curve Pro Parametric Equalizer. Furthermore, the level of sound was adjusted for each subject to ensure clear and comfortable hearing of the stories.

### MRI data collection and preprocessing

MRI data were collected on a 3T Siemens TIM Trio scanner at the Brain Imaging Center, UC Berkeley, using a 32-channel volume coil. For functional scans, a gradient echo EPI sequence was used with TR = 2.0045 s, TE = 31 ms, flip angle = 70°, voxel size = 2.24 x 2.24 x 4.1 mm^3^, matrix size = 100 x 100, field of view = 224 x 224 mm^2^ and 32 axial slices covering the entire cortex. For anatomical data, a T1-weighted multi-echo MP-RAGE sequence was used with voxel size = 1 x 1 x 1 mm^3^ and field of view = 256 x 212 x 256 mm^3^.

Each functional run was motion corrected using FMRIB’s Linear Image Registration Tool (FLIRT) (Jenkinson and Smith 2001). A cascaded motion-correction procedure was performed, where separate transformation matrices were estimated within single runs, within single sessions and across sessions sequentially. To do this, volumes in each run were realigned to the mean volume of the run. For each session, the mean volume of each run was then realigned to the mean volume of the first run in the session (see Supp. Table S1 for within-session motion statistics during the cocktail-party experiment). Lastly, the mean volume of the first run of each session was realigned to the mean volume of the first run of the first session of the passive-listening experiment. The estimated transformation matrices were concatenated and applied in a single step. Motion-corrected data were manually checked to ensure that no major realignment errors remained. The moment-to-moment variations in head position were also estimated and used as nuisance regressors during model estimation to regress out motion-related nuisance effects from BOLD responses. The Brain Extraction Tool (BET) in FSL 5.0 (Smith 2002) was used to remove non-brain tissues. This resulted in 68016-84852 brain voxels in individual subjects. All model fits and analyses were performed on these brain voxels in volumetric space.

### Visualization on cortical flatmaps

Cortical flatmaps were used for visualization purposes, where results in volumetric space were projected onto the cortical surfaces using PyCortex (Gao et al. 2015). Cortical surfaces were reconstructed from anatomical data using Freesurfer (Dale et al. 1999). Five relaxation cuts were made into the surface of each hemisphere, and the surface crossing the corpus callosum was removed. Functional data were aligned to the individual anatomical data with affine transformations using FLIRT (Jenkinson and Smith 2009). Cortical flatmaps were constructed for visualization of significant model prediction scores, functional selectivity and attentional modulation profiles, and representational complexity and modulation gradients.

### ROI definitions and abbreviations

We defined region of interests for each subject based on an atlas-based parcellation of the cortex (Destrieux et al. 2010). To do this, functional data were co-registered to the individual-subject anatomical scans with affine transformations using FLIRT (Jenkinson and Smith 2009). Individual-subject anatomical data were then registered to the Freesurfer standard anatomical space via the boundary-based registration tool in FSL (Greve and Fischl 2009). This procedure resulted in subject-specific transformations mapping between the standard anatomical space and the functional space of individual subjects. Anatomical regions of interest from the Destrieux atlas were outlined in the Freesurfer standard anatomical space; and they were back-projected onto individual-subject functional spaces via the subject-specific transformations using PyCortex (Gao et al. 2015). The anatomical regions were labeled according to the atlas. To explore potential selectivity gradients across the lateral aspects of Superior Temporal Gyrus and Superior Temporal Sulcus, these ROIs were further split into three equidistant sub-regions in posterior-to-anterior direction. Heschl’s Gyrus and Heschl’s Sulcus were considered as a single ROI as prior reports suggest that primary auditory cortex is not constrained by Heschl’s Gyrus and extends to Heschl’s Sulcus as well (Woods et al. 2009, 2010; da Costa et al. 2011). We only considered regions with at least ten speech-selective voxels in each individual subject for subsequent analyses.

Table S2 lists the defined ROIs and the number of spectrally, articulatorily and semantically selective voxels within each ROI, with number of speech-selective voxels. ROI abbreviations and corresponding Destrieux indices are: Heschl’s Gyrus and Heschl’s Sulcus (HG/HS: 33 and 74), Planum Temporale (PT: 36), posterior segment of Slyvian Fissure (pSF: 41), lateral aspect of Superior Temporal Gyrus (STG: 34), Superior Temporal Sulcus (STS, 73), Middle Temporal Gyrus (MTG: 38), Angular Gyrus (AG: 25), Supramarginal Gyrus (SMG: 26), Intraparietal Sulcus (IPS: 56), opercular part of Inferior Frontal Gyrus/Pars Opercularis (POP: 12), triangular part of Inferior Frontal Gyrus/Pars Triangularis (PTR: 14), Precentral Gyrus (PreG: 29), medial Occipito-Temporal Sulcus (mOTS:60), Inferior Frontal Sulcus (IFS: 52), Middle Frontal Gyrus (MFG:15), Middle Frontal Sulcus (MFS: 53), Superior Frontal Sulcus (SFS: 54), Superior Frontal Gyrus (SFG: 16), Precuneus (PreC: 30), Subparietal Sulcus (SPS: 71), and Posterior Cingulate Cortex (PCC: 9 and 10). The subregions of STG are: aSTG (anterior one third of STG), mSTG (middle one third of STG) and pSTG (posterior one third of STG). The subregions of STS are: aSTS (anterior one third of STS), mSTS (middle one third of STS) and pSTS (posterior one third of STS). MTG was not split into subregions since these subregions did not have a sufficient number of speech-selective voxels in each individual subject.

### Model construction

To comprehensively assess speech representations, we constructed spectral, articulatory, and semantic models of the speech stimuli (Figure 2; de Heer et al. 2017).

**Figure 2:**
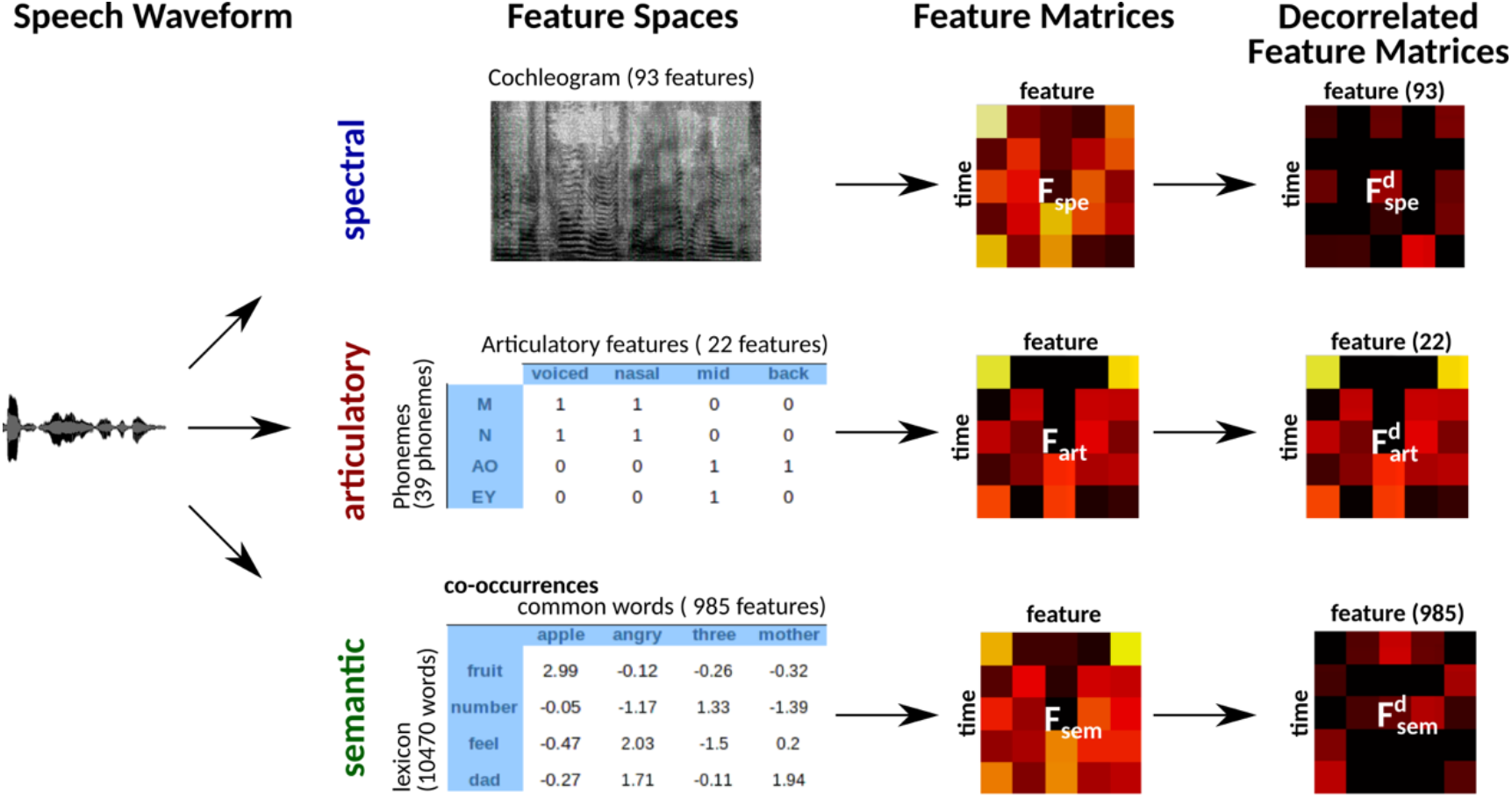
Multi-level speech features. Three distinct feature spaces were constructed to represent natural speech at multiple levels: spectral, articulatory, and semantic spaces. Speech waveforms were projected separately on these spaces to form stimulus matrices. The spectral feature matrix captured the cochleogram features of the stimulus in 93 channels having center frequencies between 115 and 9920 Hz. The articulatory feature matrix captured the mapping of each phoneme in the stimulus to 22 binary articulation features. The semantic feature matrix captured the statistical co-occurrences of each word in the stimulus with 985 common words in English. Each feature matrix was Lanczos-filtered at a cutoff frequency of 0.25 Hz and downsampled to 0.5 Hz to match the sampling rate of fMRI. Natural speech might contain intrinsic stimulus correlations among spectral, articulatory, and semantic features. To prevent potential biases due to stimulus correlations, we decorrelated the three feature matrices examined here via Gram-Schmidt orthogonalization (see Methods). The decorrelated feature matrices were used for modeling BOLD responses.

#### Spectral model

For the spectral model, cochleogram features of speech were estimated based on Lyon’s Passive Ear model. Lyon’s human cochlear model involves logarithmic filtering, compression and adaptive gain control operations applied to input sound (Lyon 1982; Slaney 1998; Gill et al. 2006). Depending on the sampling rate of the input signal, the cochlear model generates 118 waveforms with center frequencies between ∼84 Hz and ∼21 kHz. Considering the frequency response of the headphones used in the experiment, 93 waveforms with center frequencies between 115 Hz and 9920 Hz were selected as the features of the spectral model. The spectral features were Lanczos-filtered at a cutoff frequency of 0.25 Hz and downsampled to 0.5 Hz to match the sampling rate of fMRI. The 93 spectral features were then temporally z-scored to zero mean and unit variance.

#### Articulatory model

For the articulatory model, each phoneme in the stories was mapped onto a unique set of 22 articulation features; for example, phoneme /ʒ/ is postalveolar, fricative and voiced (Levelt 1993; de Heer et al. 2017). This mapping resulted in 22-dimensional binary vectors for each phoneme. To obtain the timestamp of each phoneme and word in the stimuli, the speech in the stories were aligned with the story transcriptions using the Penn Phonetics Lab Forced Aligner (Yuan and Liberman 2008). Alignments were manually verified and corrected using Praat (www.praat.org). The articulatory features were Lanczos-filtered at a cutoff frequency of 0.25 Hz and downsampled to 0.5 Hz. Finally, the 22 articulatory features were z-scored to zero mean and unit variance.

#### Semantic model

For the semantic model, co-occurrence statistics of words were measured via a large corpus of text (Mitchell et al. 2008; Huth et al. 2016; de Heer et al. 2017). The text corpus was compiled from 2,405,569 Wikipedia pages, 36,333,459 user comments scraped from reddit.com, 604 popular books and the transcripts of 13 Moth stories (including the stories used as stimuli). We then built a 10,470-word lexicon from the union set of the 10,000 most common words in the compiled corpus and all words appearing in the ten Moth stories used in the experiment. Basis words were then selected as a set of 985 unique words from Wikipedia’s List of 1000 Basic Words. Co-occurrence statistics of the lexicon words with 985 basis words within a 15-word window were characterized as a co-occurrence matrix of size 985×10,470. Elements of the resulting co-occurrence matrix were log-transformed, z-scored across columns to correct for differences in basis-word frequency, and z-scored across rows to correct for differences in lexicon-word frequency. Each word in the stimuli was then represented with a 985-dimensional co-occurrence vector based on the speech-transcription alignments. The semantic features were Lanczos-filtered at a cutoff frequency of 0.25 Hz and downsampled to 0.5 Hz. The 985 semantic features were finally z-scored to zero mean and unit variance.

#### Decorrelation of feature spaces

In natural stories, there might be potential correlations among certain spectral, articulatory, or semantic features. If significant, such correlations can partly confound assessments of model performance. To assess the unique contribution of each feature space to the explained variance in BOLD responses, a decorrelation procedure was first performed (Figure 2). To decorrelate a feature matrix F of size *mxn* from a second feature matrix K of size *mxp*, we first found an orthonormal basis for the column space of K (*col*{*K*}) using economy-size singular value decomposition:

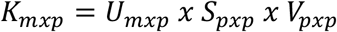

where U contains left singular vectors as columns, V contains right singular vectors, and S contains the singular values. Left singular vectors were taken as the orthonormal basis for col{K}, and each column of F was decorrelated from it according to the following formula:

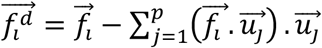

Where 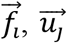 are the column vectors of F and U respectively, and 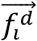 is the column vectors of the decorrelated feature matrix, *F^d^*. To decorrelate feature matrices for the models considered here, we took the original articulatory feature matrix as a reference, and decorrelated the spectral feature matrix from the articulatory feature matrix, and decorrelated the semantic feature matrix from both articulatory and spectral feature matrices. This decorrelation sequence was selected because spectral and articulatory features capture lower-level speech representations, and the articulatory feature matrix had the fewest number of features among all models. In the end, we obtained 3 decorrelated feature matrices whose columns had zero correlation with the columns of the other two matrices.

### Analyses

The main motivation of this study is to understand whether and how strongly various levels of speech representations are modulated across cortex during a cocktail-party task. To answer this question, we followed a two-stage approach as illustrated in Figure 3. In the first stage, we identified voxels selective for speech features using data from the passive-listening experiment. To do this, we measured voxelwise selectivity separately for spectral, articulatory, and semantic features of the single-speaker stories. In the second stage, we used the models fit using passive-listening data to predict BOLD responses measured in the cocktail-party experiment. Prediction scores for attended versus unattended stories were compared to quantify the degree of attentional modulations, separately for each model and globally across all models.

**Figure 3:**
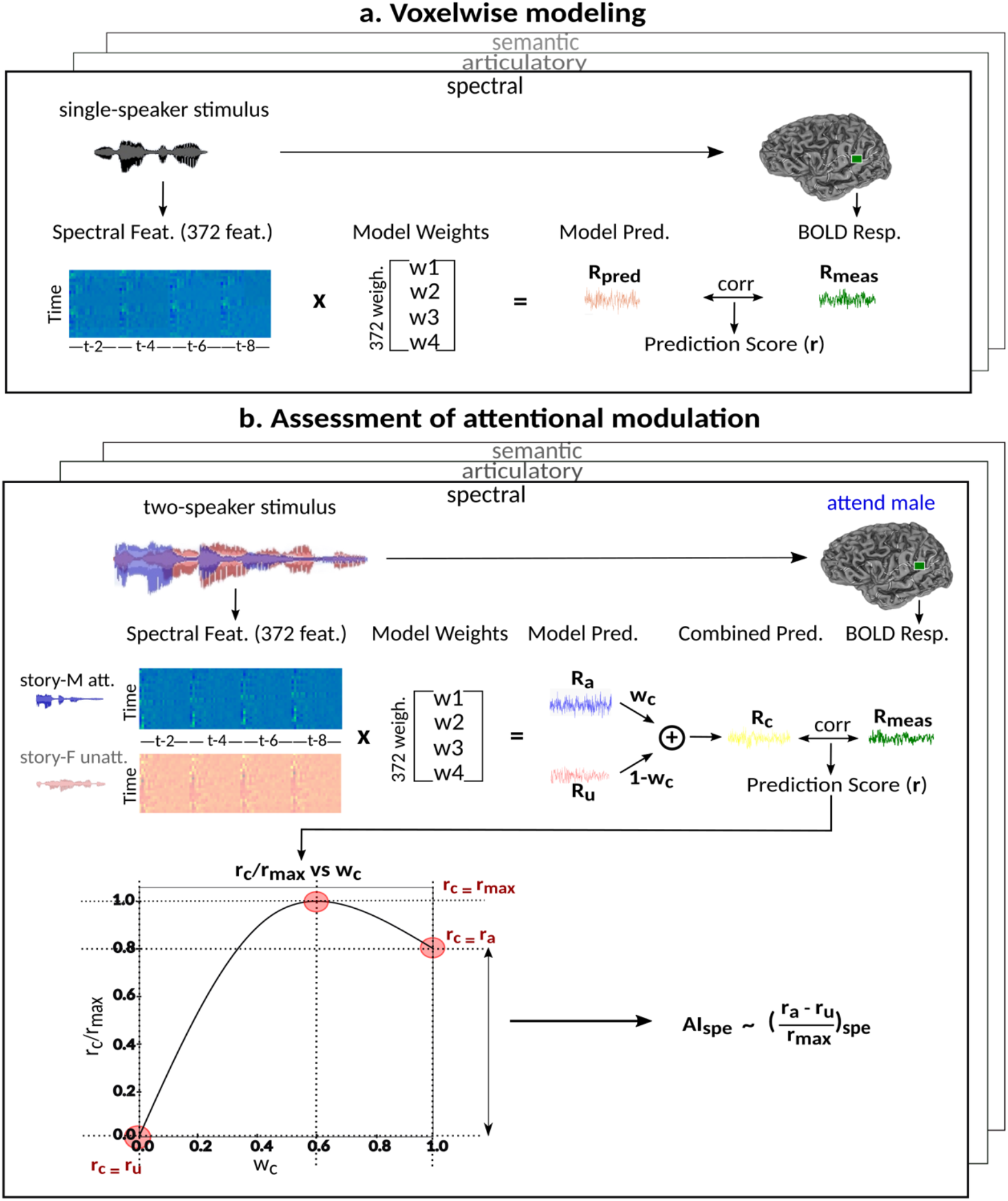
Modeling procedures. **a.** *Voxelwise modeling.* Voxelwise models were fit in individual subjects using passive-listening data. To account for hemodynamic response, a linearized four-tap finite impulse response (FIR) filter spanning delayed effects at 2-8 sec was used. Models were fit via L2-regularized linear regression. BOLD responses were predicted based on fit voxelwise models on held-out passive-listening data. Prediction scores were taken as the Pearson’s correlation between predicted and measured BOLD responses. For a given subject, speech-selective voxels were taken as the union of voxels significantly predicted by spectral, articulatory, or semantic models (q(FDR) < 10^-5^, t-test). **b.** *Assessment of attentional modulation.* Passive-listening models for single voxels were tested on cocktail-party data to quantify attentional modulations in selectivity. In a given run, one of the speakers in a two-speaker story was attended while the other speaker was ignored. Separate response predictions were obtained using the isolated story stimuli for the attended speaker and for the unattended speaker. Since a voxel can represent information from both attended and unattended stimuli, a linear combination of these predicted responses was considered with varying combination weights (wc in [0 1]). BOLD responses were predicted based on each combination weight separately. Three separate prediction scores were calculated based on only the attended stimulus (wc=1), based on only the unattended stimulus (wc=0), and based on the optimal combination of the two stimuli. A model-specific attention index, (*AI_m_*) was then computed as the ratio of the difference in prediction scores for attended versus unattended stories to the prediction score for their optimal combination (see Methods).

Note that a subset of the ten single-speaker-stories were used to generate three two-speaker-stories used in the experiments. To prevent potential bias, a three-fold cross-validation procedure was performed for testing models fit using passive-listening data on cocktail-party data. In each fold, models were fit using eight-run passive-listening data; and separately tested on two-run passive-listening data and two-run cocktail-party data. The same set of test stories were used both in the passive-listening and cocktail-party experiments to minimize risk of poor model generalization between the passive-listening and cocktail-party experiments due to uncontrolled stimulus differences. There was no overlap between the stories in the training and testing runs. Model predictions were aggregated across three folds, and prediction scores were then computed.

#### Voxelwise modeling

In the first stage, we fit voxelwise models in individual subjects using passive-listening data. To account for hemodynamic delays, we used a linearized four-tap finite impulse response (FIR) filter to allow different HRF shapes for separate brain regions (Goutte et al. 2000). Each model feature was represented as four features in the stimulus matrix to account for their delayed effects in BOLD responses at 2, 4, 6 and 8 sec. Model weights, *W*, were then found using L2-regularized linear regression:

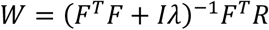

Here, *λ* is the regularization parameter, *F* is the decorrelated feature matrix for a given model and *R* is the aggregate BOLD response matrix for cortical voxels. A cross-validation procedure with 50 iterations was performed to find the best regularization parameter for each voxel among 30 equispaced values in log-space of 1: 10^5^. The training passive-listening data was split into 50 equisized chunks, where 1 chunk was reserved for validation and 49 chunks were reserved for model fitting at each iteration. Prediction scores were taken as Pearson’s correlation between predicted and measured BOLD responses. The optimal *λ* value for each voxel was selected by maximizing the average prediction score across cross-validation folds. The final model weights were obtained using the entire set of training passive-listening data and the optimal *λ*. Next, we measured the prediction scores of the fit models on testing data from the passive-listening experiment. Spectrally-, articulatorily-, and semantically-selective voxels were separately identified in each ROI based on the set of significantly-predicted voxels by each model. A given ROI was considered selective for a model, only if it contained ten or more significant voxels for that model (*q*(*FDR*) < 10^-5^; *t*; − *t*;*est*;). Speech-selective voxels within the ROI were then taken as the union of these spectrally-, articulatorily-, and semantically-selective voxels. Subsequent analyses were performed on speech-selective voxels.

a. **Model-specific selectivity index.** Single-voxel prediction scores on passive-listening data were used to quantify the degree of selectivity of each ROI to the underlying model features under passive-listening. To do this, a model-specific selectivity index, (*SI_m_*), was defined as follows:

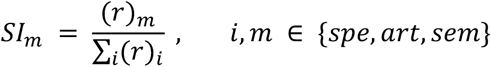

where *r* is the average prediction score across speech-selective voxels within the ROI during passive-listening. *SI_m_* is in the range of [0, 1], where higher values indicate stronger selectivity for the underlying model.
b. **Complexity index.** The complexity of speech representations was characterized via a complexity index, (*CI*), which reflected the relative tuning of an ROI for low-versus high-level speech features. The following intrinsic complexity levels were assumed for the three speech models considered here: (*c_spe_*, *c_art_*, *c_sem_*) = (0.0, 0.5, 1.0). Afterwards, *CI* was taken as the average of the complexity levels weighted by the selectivity indices:

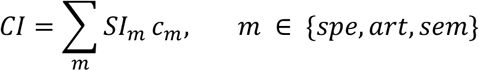

*CI* is in the range of [0, 1], where higher values indicate stronger tuning for semantic features and lower values indicate stronger tuning for spectral features.

#### Assessment of attentional modulations

In the second stage, we tested the passive-listening models on cocktail-party data to quantify ROI-wise attentional modulation in selectivity for corresponding model features and to find the extent of the representation of unattended speech. These analyses were repeated separately for the three speech models.

a. **Model-specific attention index.** To quantify the attentional modulation in selectivity for speech features, we compared prediction scores for attended versus unattended stories in the cocktail-party experiment. Models fit using passive-listening data were used to predict BOLD responses elicited by two-speaker stories. In each run, only one of the speakers in a two-speaker story was attended while the other speaker was ignored. Separate response predictions were obtained using the isolated story stimuli for the attended and unattended speakers. Since a voxel can represent information on both attended and unattended stimuli, a weighted linear combination of these predicted responses was considered:

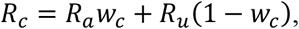

Where *R_a_* and *R_u_* are the predicted responses for the attended and unattended stories in a given run; *R_c_* is the combined response and *w_c_* is the combination weight. We computed *R_c_* for each separate *w_c_* value in [0:0.1:1]. Note that *R_c_* = *R_a_* when *w_c_* = 1.0; and *R_c_* = *R_u_* when *w_c_* = 0.0. We then calculated single-voxel prediction scores for each *w_e_* value. An illustrative plot of *r_c_*⁄*r_max_* vs *w_c_* is given in Figure 3b, where *r_c_* denotes the prediction scores and *r_max_* denotes the maximum *r_c_* value (the optimal combination). *r_a_* and *r_u_* are the prediction scores for attended and unattended stories respectively. To quantify the degree of attentional modulation, a model-specific attention index (*AI_m_*) was taken as:

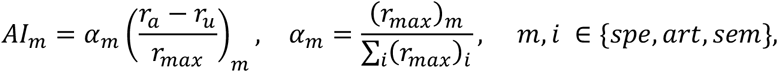

where *r_max_* denotes an ideal upper limit for model performance, and *α_m_* reflects the relative model performance under the cocktail-party task. Note that *AI_m_* considers selectivity to the underlying model features when calculating the degree of attentional modulation.
b. **Global attention index.** We then computed global attention index (*gAI*) as follows:

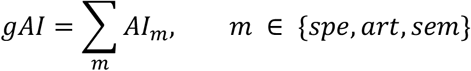

Both *gAI* and *AI_m_* are in the range [-1,1]. A positive index indicates attentional modulation of selectivity in favor of the attended stimuli and a negative index indicates attentional modulation in favor of the unattended stimuli. A value of zero indicates no modulation.

#### Colormap in selectivity and modulation profile flatmaps

The cortical flatmaps of selectivity and modulation profiles use a colormap that shows the relative contributions of all three models to the selectivity and attention profiles. For selectivity profiles, a continuous colormap was created by assigning significantly positive articulatory, semantic and spectral selectivity to the red, green and blue (R, G, B) color channels, respectively. During assignment, selectivity values were normalized to sum of one, and then normalized to linearly map the interval [0.15 0.85] to [0 1]. Distinct colors were assigned to six landmark selectivity values: red for (1, 0, 0), green for (0,1,0), blue for (0,0, 1), yellow for (0.5,0.5,0), magenta for (0.5, 0, 0.5), and turquoise for (0, 0.5, 0.5). The same procedures were also applied for creating a colormap for modulation profiles.

### Statistical Tests

#### Significance assessments within subjects

For each voxel-wise model, significance of prediction scores was assessed via a t-test; and resulting p-values were false-discovery-rate corrected for multiple comparisons (FDR; Benjamini and Hochberg 1995).

A bootstrap test was used in assessments of *SI_m_*, *CI*, *AI_m_* and *gAI* within single subjects. In ROI analyses, speech-selective voxels within a given ROI were resampled with replacement 10000 times. For each bootstrap sample, mean prediction score of a given model was computed across resampled voxels. Significance level was taken as the fraction of bootstrap samples in which the test metric computed from these prediction scores is less than 0 (for right-sided tests) or greater than 0 (for left-sided tests). The same procedure was also used for comparing pairs of ROIs, where ROI voxels were resampled independently.

#### Significance assessments across subjects

A bootstrap test was used in assessments of *SI_m_*, *CI*, *AI_m_* and *gAI* across subjects. In ROI analyses, ROI-wise metrics were resampled across subjects with replacement 10000 times. Significance level was taken as the fraction of bootstrap samples where the test metric averaged across resampled subjects is less than 0 (for right-sided tests) or greater than 0 (for left-sided tests). The same procedure was also used for comparisons among pairs of ROIs. Here, we used a more stringent significance definition for across-subjects tests that focuses on effects consistently observed in each individual subject. Therefore, an effect was taken significant only if the same metric was found significant in each individual subject.

## Results

### Attentional modulation of multi-level speech representations

To examine the cortical distribution and strength of attention modulations in speech representations, we first obtained a baseline measure of intrinsic selectivity for speech features. For this purpose, we fit voxelwise models using BOLD responses recorded during passive listening. We built three separate models containing low-level spectral, intermediate-level articulatory and high-level semantic features of natural stories (de Heer et al. 2017). Supplementary Fig. S1 displays the cortical distribution of prediction scores for each model in a representative subject, and Supp. Table S2 lists the number of significantly predicted voxels by each model in anatomical ROIs. We find *spectrally-selective voxels* mainly in early auditory regions (bilateral HG/HS and PT; and left pSF) and bilateral SMG, and a*rticulatorily-selective voxels* mainly in early auditory regions (bilateral HG/HS and PT; and left pSF), bilateral STG, STS, SMG and MFS as well as left POP and PreG. In contrast, s*emantically-selective voxels* are found broadly across cortex except early auditory regions (bilateral HG/HS and right PT).

To quantitatively examine cortical overlap among spectral, articulatory, and semantic representations, we separately measured the degree of functional selectivity for each feature level via a model-specific selectivity index (*SI_m_*; see Methods). Bar plots of selectivity indices are displayed in Figure 4a for perisylvian cortex and in Supplementary Fig. S2 for non-perisylvian cortex (see Supp. Fig. S3a-e for single-subject results). Distinct selectivity profiles are observed from distributed selectivity for spectral, articulatory, and semantic features (e.g., left PT and right pSTG) to strong tuning to a single level of features (e.g., left IPS and PCC). The selectivity profiles of the ROIs are also visualized on the cortical flatmap projecting articulatory, semantic, and spectral selectivity indices of each ROI to the red, green, and blue channels of the RGB colormap as seen in Figure 4b (see Supplementary Fig. S4 for selectivity profile flatmaps in individual subjects; see Methods for colormap details). A progression from low-intermediate to high-level speech representations is apparent across bilateral temporal cortex in superior-inferior direction (HG/HS → mSTG → mSTS → MTG) consistently in all subjects. Furthermore, many higher-order regions in parietal (bilateral AG, IPS, SPS, PrC, PCC and POS) and frontal cortices (bilateral PTR, IFS, MFG, SFS, and SFG; and left POP) manifest dominant semantic selectivity consistently in all subjects (p < 0.05; see Supp. Fig. S3a-e for single-subject results). To examine the hierarchical organization of the speech representations in a finer scale, we also defined a complexity index, *CI*, that reflects whether an ROI is relatively tuned for low-level spectral or high-level semantic features. A detailed investigation of the gradients in *CI* across two main auditory streams (dorsal and ventral stream) was conducted (see Supplementary Results). These results corroborate the view that speech representations are hierarchically organized across cortex with partial overlap mostly in early and intermediate stages of speech processing.

**Figure 4:**
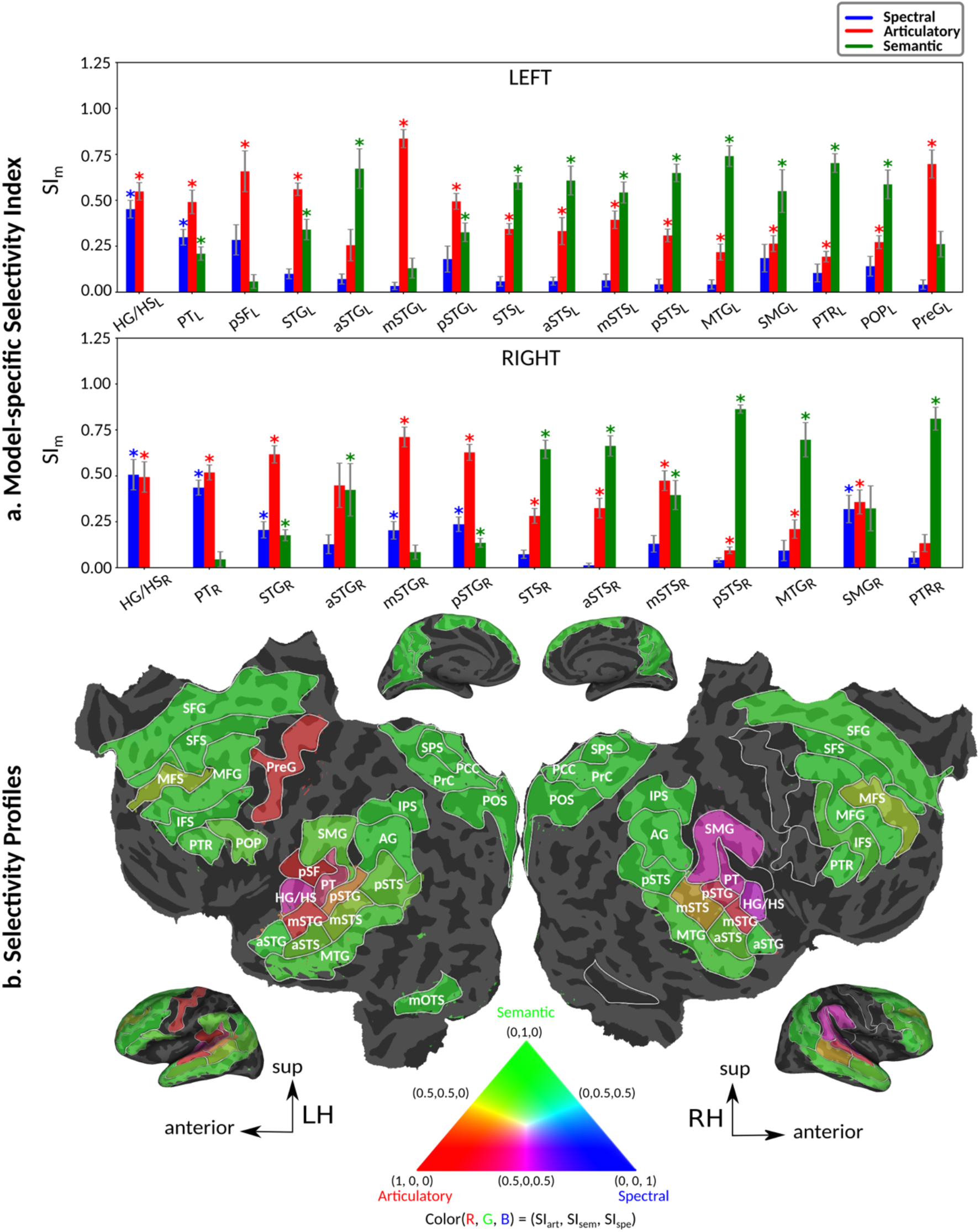
Selectivity for multi-level speech features. **a.** *Model-specific selectivity indices*. Single-voxel prediction scores on passive-listening data were used to quantify the selectivity of each ROI to underlying model features. Model-specific prediction scores were averaged across speech-selective voxels within each ROI and normalized such that the cumulative score from all models was 1. The resultant measure was taken as a model-specific selectivity index, (*SI_m_*). *SI_m_* is in the range of [0, 1], where higher values indicate stronger selectivity for the underlying model. Bar plots display *SI_m_* for spectral, articulatory, and semantic models (mean ± sem across subjects). Significant indices are marked with * (p < 0.05; see Sup. Fig. S3a-e for selectivity indices of individual subjects). ROIs in perisylvian cortex are displayed (see Supp. Fig. S2 for non-perisylvian ROIs; see Methods for ROI abbreviations). ROIs in LH and RH are shown in the top and bottom panels, respectively. POPR and PreGR that did not have consistent speech selectivity in individual subjects were excluded (see Methods). **b.** *Intrinsic selectivity profiles.* Selectivity profiles of cortical ROIs averaged across subjects are shown on the cortical flatmap of a representative subject (S4). Significant articulatory, semantic, and spectral selectivity indices of each ROI are projected to the red, green, and blue channels of the RGB colormap (see Methods). This analysis only included ROIs with consistent selectivity for speech features in each individual subject. Medial and lateral views of the inflated hemispheres are also shown. A progression from low-intermediate to high-level speech representations are apparent across bilateral temporal cortex in the superior-inferior direction; consistently in all subjects (see Supp. Fig. S4 for selectivity profiles of individual subjects). Meanwhile, semantic selectivity is dominant in many higher-order regions within the parietal and frontal cortices (bilateral AG, IPS, SPS, PrC, PCC, POS, PTR, IFS, SFS, SFG, MFG and left POP) (p < 0.05; see Supp. Fig. S3a-e). These results support the view that speech representations are hierarchically organized across cortex with partial overlap between spectral, articulatory and semantic representations in early to intermediate stages of auditory processing.

Next, we systematically examined attentional modulations at each level of speech representation during a diotic cocktail-party task. To do this, we recorded whole-brain BOLD responses while participants listened to temporally-overlaid spoken narratives from two different speakers and attended to either a male or female speaker in these two-speaker stories. We used the spectral, articulatory, and semantic models fit using passive-listening data to predict responses during the cocktail-party task. Since a voxel can represent information on both attended and unattended stimuli, response predictions were expressed as a convex combination of individual predictions for the attended and unattended story within each two-speaker story. Prediction scores were computed based on estimated responses as the combination weights were varied in [0 1] (see Methods). Scores for the optimal combination model were compared against the scores from the individual models for attended and unattended stories. If the optimal combination model significantly outperforms the individual models, it indicates that the voxel represents information from both attended and unattended stimuli.

Figure 5 displays prediction scores of the spectral, articulatory, and semantic models as a function of the combination weight in representative ROIs, including HG, HS and PT. Scores based on only attended story (*r_a_*), based on only the unattended story (*r_u_*), and based on the optimal combination of the two (*r_max_*) are marked. A diverse set of attentional effects are observed for each type of model. For the *spectral model* in left HG/HS, the optimal combination assigns matched weights to attended and unattended stories, and *r_max_* is larger than *r_a_* (p < 10^-4^). This finding implies that spectral representations of the unattended story are mostly maintained; and there is no apparent bias towards the attended story at spectral level in left HG/HS. For the *articulatory model* in left HG/HS, *r_a_* is larger than *r_u_* (p < 10^-4^), while *r_max_* is greater than *r_a_* (p < 10^-2^). Besides, the optimal combination gives slightly higher weight to the attended versus unattended story. This result suggests that attention moderately shifts articulatory representations in left HG/HS in favor of the attended stream such that articulatory representations of the unattended story are preserved to a degree. For the s*emantic model* in left PT, *r_a_* is much higher than *r_u_* (p < 10^-4^). Besides, the optimal combination assigns substantially higher weight to the attended story in this case. This finding indicates that attention strongly shifts semantic representations in left PT towards the attended stimulus. A simple inspection of these results suggests that attention may have distinct effects at various levels of speech representation across cortex. Hence, a detailed quantitative analysis is warranted to measure the effect of attention at each level.

**Figure 5:**
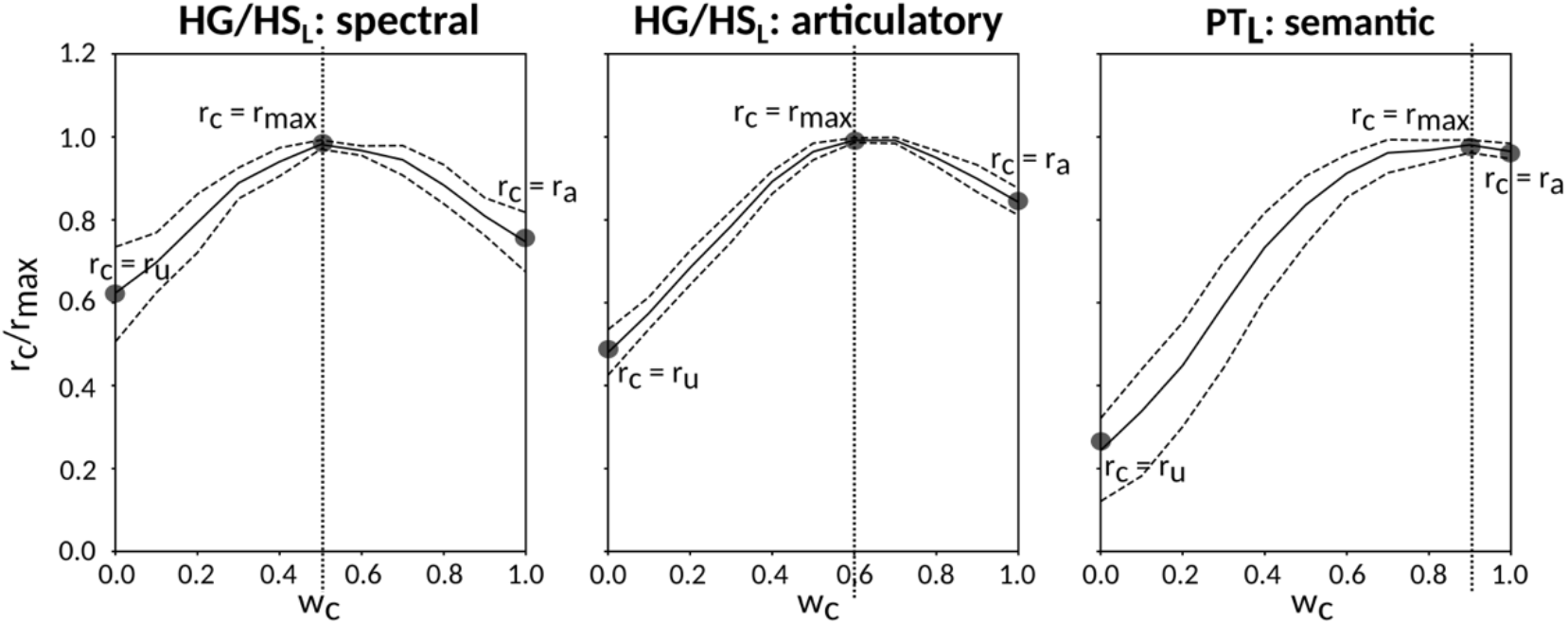
Predicting cocktail-party responses. Passive-listening models were tested during the cocktail-party task by predicting BOLD responses in the cocktail-party data. Since a voxel might represent information from both attended and unattended stimuli, response predictions were expressed as a convex combination of individual predictions for the attended and unattended story within each two-speaker story. Prediction scores were computed as the combination weights (*w_c_*) were varied in [0 1] (see Methods). Prediction scores for a given model were averaged across speech-selective voxels within each ROI (*r_c_*). The normalized scores of spectral, articulatory, and semantic models are displayed in several representative ROIs (HG/HS, HG/HS, and PT). Solid and dashed lines indicate mean and %95 confidence intervals across subjects. Scores based on only the attended story (*r_a_*), based on only the unattended story (*r_u_*), and based on the optimal combination of the two (*r_max_*) are marked with circles. For the *spectral model* in left HG/HS, *r_max_* is larger than *r_a_* (p < 10^-4^); and the optimal combination equally weighs attended and unattended stories. For the *articulatory model* in left HG/HS, *r_a_* is larger than *r_u_* (p < 10^-4^), while *r_max_* is greater than *r_a_* (p < 10^-2^). Besides, the optimal combination puts slightly higher weight to attended story than unattended story. For the s*emantic model* in left PT, *r_a_* is much higher than *r_u_* (p < 10^-4^), and the optimal combination puts much greater weight to attended story than unattended one. These representative results imply that attention may have divergent effects at various levels of speech representations across cortex.

#### Level-specific attentional modulations

To quantitatively assess the strength and direction of attentional modulations, we separately investigated the modulatory effects on spectral, articulatory, and semantic features across cortex. To measure modulatory effects at each feature level, a model-specific attention index (*AI_m_*) was computed, reflecting the difference in model prediction scores when the stories were attended versus unattended (see Methods). *AI_m_* is in the range of [-1, 1]; a positive index indicates selectivity modulation in favor of the attended stimulus, whereas a negative index indicates selectivity modulation in favor of the unattended stimulus. A value of zero indicates no modulation.

Figure 6a and Supp. Fig. S7 display the attention index for spectral, articulatory, and semantic models across perisylvian and non-perislyvian ROIs, respectively (see Supp. Fig. S8a-e for single-subject results). The modulation profiles of the ROIs are also visualized on the cortical flatmap, projecting articulatory, semantic, and spectral attention indices to the red, green, and blue channels of the RGB colormap as seen in Figure 6b (see Supp. Fig. S9 for modulation profile flatmaps in individual subjects). Here we discuss the attention index for each model individually. *Spectral modulation* is not consistently significant in each subject across perisylvian ROIs (p > 0.05). On the other hand, moderate spectral modulation is found in right SFG consistently in all subjects (p < 10^-3^). *Articulatory modulation* starts as early as HG/HS bilaterally (p < 10^-3^). In the dorsal stream, it extends to PreG and POP in the left hemisphere (LH) and to SMG in the right hemisphere (RH; p < 10^-2^); and it becomes dominant only in left PreG consistently in all subjects (p < 0.05). In the ventral stream, it extends to left PTR and bilateral MTG (p < 10^-2^). Articulatory modulation is also found -albeit generally less strongly- in frontal regions (bilateral MFS; left MFG; and right IFS and SFG) consistently in all subjects (p < 0.05). In the dorsal stream, *semantic modulation* starts in PT and extends to POP in LH (p < 10^-2^), whereas it is not apparent in the right dorsal stream (p > 0.05). In the ventral stream, semantic modulation starts in aSTG and mSTS bilaterally (p < 0.05). It extends to MTG and PTR, and becomes dominant in both ends of the bilateral ventral stream (p < 0.05). Lastly, semantic modulation is observed widespread across higher-order regions within frontal and parietal cortices consistently in all subjects (p < 0.05), with the exception of left IPS (p > 0.05). Taken together, these results suggest that attending to a target speaker alters articulatory and semantic representations broadly across cortex.

**Figure 6:**
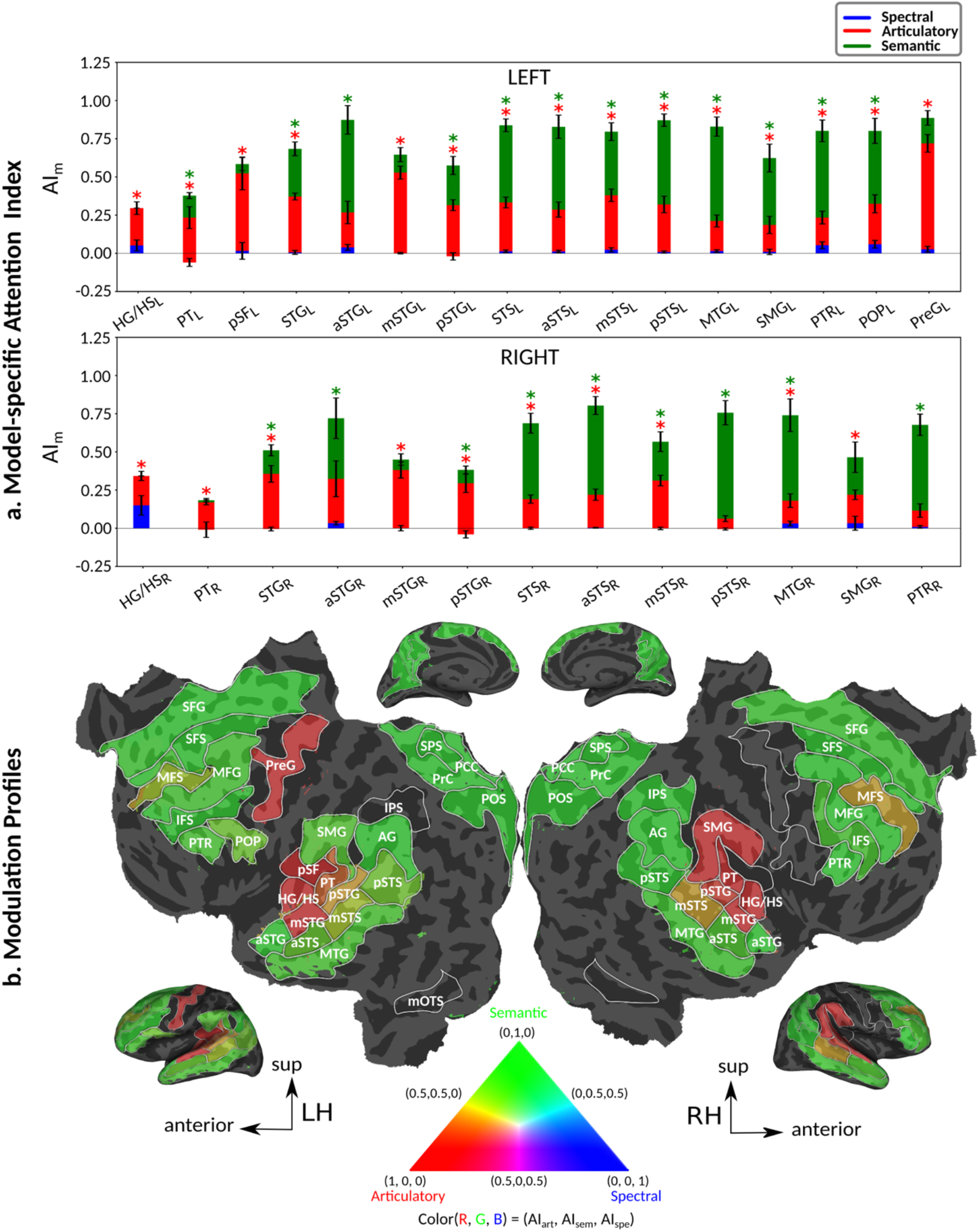
Attentional modulation of multi-level speech representations. **a.** *Model-specific attention indices.* A model-specific attention index (*AI_m_*) was computed based on the difference in model prediction scores when the stories were attended versus unattended (see Methods). *AI_m_* is in the range of [-1,1], where a positive index indicates modulation in favor of the attended stimulus and a negative index indicates modulation in favor of the unattended stimulus. For each ROI in perisylvian cortex, spectral, articulatory, and semantic attention indices are given (mean ± sem across subjects), and their sum yields the overall modulation (see Supp. Fig. S7 for non-perisylvian ROIs). Significantly positive indices are marked with * (p < 0.05, bootstrap test; see Supp. Fig. S8a-e for attention indices of individual subjects). ROIs in the LH and RH are shown in top and bottom panels, respectively. These results show that selectivity modulations distribute broadly across cortex at the linguistic level (articulatory and semantic). **b.** *Attentional modulation profiles.* Modulation profiles averaged across subjects are displayed on the flattened cortical surface of a representative subject (S4). Significantly positive articulatory, semantic, and spectral attention indices are projected onto the red, green and blue channels of the colormap (see Methods). A progression in the level of speech representations dominantly modulated is apparent from HG/HS to MTG across bilateral temporal cortex (see Supp. Fig. S9 for modulation profiles of individual subjects). Articulatory modulation is dominant in one end of the dorsal stream (left PreG), whereas semantic modulation becomes dominant in both ends of the ventral stream (bilateral PTR and MTG) (p < 0.05; see Supp. Figs. S8a-e and S9). On the other hand, semantic modulation is dominant in most of the higher-order regions in the parietal and frontal cortices consistently in all subjects (bilateral AG, SPS, PrC, PCC, POS, SFG, SFS, and PTR; left MFG; and right IPS) (p < 0.05; see Supp. Fig. S8a-e).

#### Global attentional modulations

It is commonly assumed that attentional effects grow stronger towards higher-order regions across the cortical hierarchy of speech (Zion Golumbic et al. 2013; O’Sullivan et al. 2019; Regev et al. 2019). Yet, a systematic examination of attentional modulation gradients across dorsal and ventral streams is lacking. To examine this issue, we measured overall attentional modulation in each region via a global attention index (*gAI*; see Methods). Similar to the model-specific attention indices, a positive *gAI* indicates modulations in favor of the attended stimulus, and a negative *gAI* indicates modulations in favor of the unattended stimulus.

a. **Dorsal stream.** We first examined variation of *gAI* across the dorsal stream (left dorsal-1: HG/HS_L_ → PT_L_ → (SMG_L_) → POP_L_, left dorsal-2: HG/HS_L_ → PT_L_ → (SMG_L_) → PreG_L_, and right dorsal: HG/HS_R_ →PT_R_ → SMG_R_) as shown in Figure 7. We find significant increase in *gAI* across the following left dorsal subtrajectories consistently in all subjects (p < 0.05; see Supp. Fig. S11 for gradients in individual subjects): *gAI_PT_* < *gAI_SMG_* < *gAI_POP_* and *gAI_PT_* < *gAI_SMG_* < *gAI_PreG_*. In contrast, we find no consistent gradient in the right dorsal stream (p > 0.05). These results suggest that attentional modulations grow progressively stronger across the dorsal stream in LH.
b. **Ventral stream.** We then examined variation of *gAI* across the ventral stream (left ventral-1: HG/HS_L_ → mSTG_L_ → mSTS_L_ → MTG_L_, left ventral-2: HG/HS_L_ → mSTG_L_ → aSTG_L_ → PTR_L_, right ventral-1: HG/HS_R_ → mSTG_R_ → mSTS_R_ → MTG_R_ and right ventral-2: HG/HS_R_ → mSTG_R_ → aSTG_R_ → PTR_R_), as shown in Figure 7. We find significant increase in *gAI* across the following subtrajectories consistently in all subjects (p < 0.05; see Supp. Fig. S11 for gradients in individual subjects): *gAI_HG/HS_* < *gAI_mSTG_* < *gAI_PSTG_* and *gAI_HG/HS_* < *gAI_mSTG_* < *gAI_mSTS_* in the left ventral stream, and *gAI_mSTG_* < *gAI_aSTG_* in the right ventral stream. In contrast, we find no difference between aSTG and PTR bilaterally, between mSTS and MTG in the left ventral stream, and between HG/HS, mSTG, mSTS and MTG in the right ventral stream (p > 0.05). These results suggest that overall attentional modulations gradually increase across the ventral stream, and that the increases are more consistent in LH compared to RH.
c. **Representational complexity versus attentional modulation.** Visual inspection of Supp. Fig. S6b and Figure 7b suggests that the subtrajectories with significant increases in *CI* and in *gAI* overlap largely in left ventral stream and partly in left dorsal stream. To quantitatively examine the overlap in left ventral stream, we analyzed the correlation between CI and gAI across the left-ventral subtrajectories where significant increases in *CI* are observed. We find significant correlations in *HG*/*HS* → *mSTG* → *mSTS* and in *HG*/*HS* → *mSTG* → *aSTG* (r > 0.98, bootstrap test, p < 10^-4^) consistently in all subjects. In line with a recent study arguing for stronger attentional modulation and higher representational complexity in STG compared to HG (O’Sullivan et al. 2019), our results indicate that attentional modulation increases towards higher-order regions as the representational complexity increases across the dorsal and ventral streams in LH (more apparent in ventral than dorsal stream).
d. **Hemispheric asymmetries in attentional modulation.** To assess potential hemispheric asymmetries in attentional modulation, we compared *gAI* between the left and right hemispheric counterparts of each ROI. This analysis was restricted to ROIs with consistent selectivity for speech features in both hemispheres in each individual subject (see Methods). Supplementary Table S3 lists the results of the across-hemisphere comparison. No consistent hemispheric asymmetry is found across cortex with the exception of mSTG having a left-hemispheric bias in *gAI* consistently in all subjects (p < 0.05). These results indicate that there is mild lateralization in attentional modulation of intermediate-level speech features.

**Figure 7:**
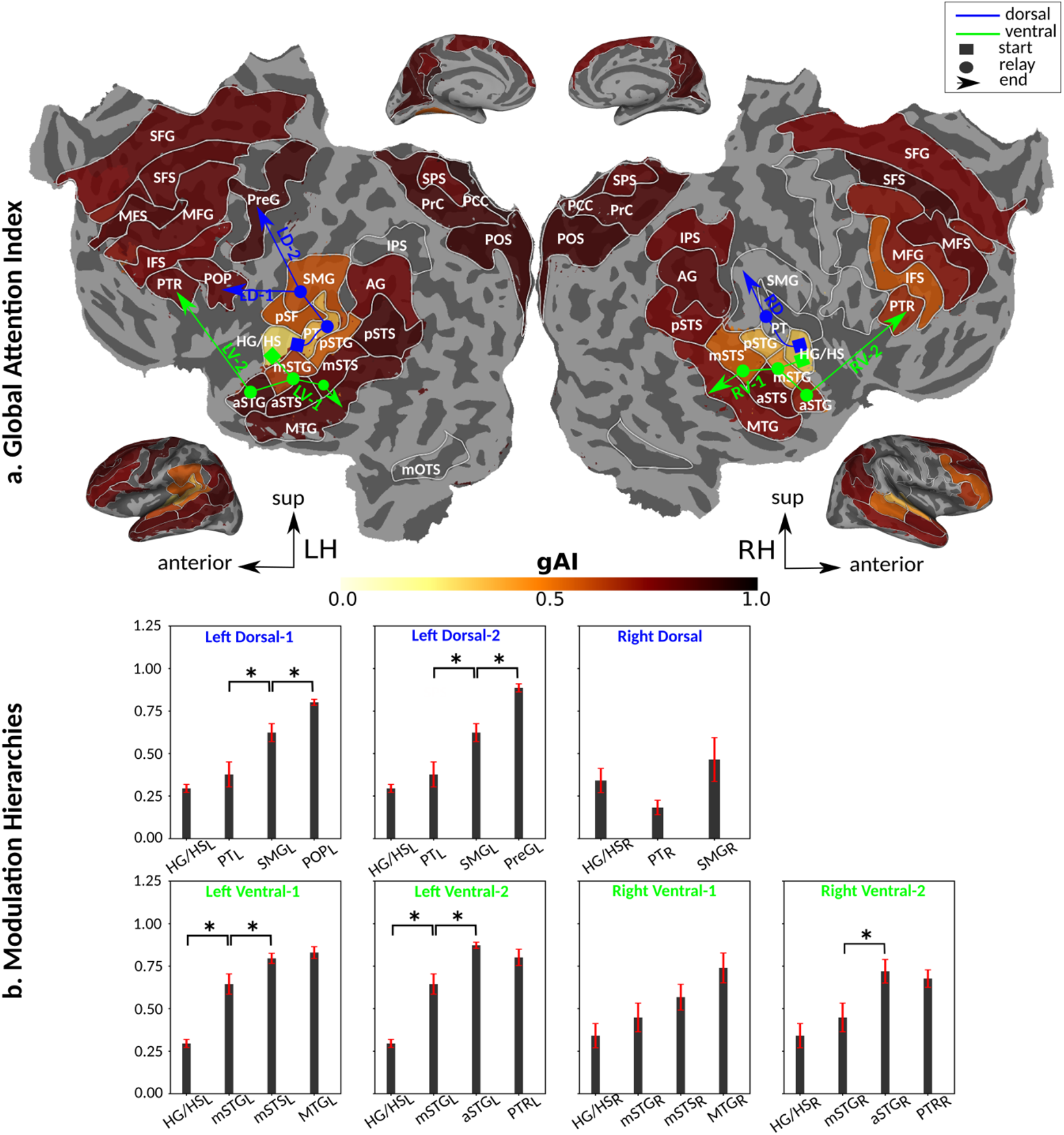
Global attentional modulation. **a.** *Global attention index.* To quantify overall modulatory effects on selectivity across all examined feature levels, global attentional modulation (*gAI*) was computed by summing spectral, articulatory, and semantic attention indices (see Methods). *gAI* is in the range of [-1,1] and a value of zero indicates no modulation. Colors indicate significantly positive *gAI* averaged across subjects (see legend; see Supp. Fig. S10 for bar plots of *gAI* across cortex). Dorsal and ventral pathways are shown with blue and green lines, respectively: left dorsal-1 (LD-1), left dorsal-2 (LD-2) and right dorsal (RD), left ventral-1 (LV-1), left ventral-2 (LV-2), right ventral-1 (RV-1) and right ventral-2 (RV-2). Squares mark regions where pathways begin; arrows mark regions where pathways end; and circles mark relay regions in between. **b.** *Modulation hierarchies.* Bar plots display *gAI* (mean ± sem across subjects) along LD-1, LD-2, RD, LV-1, LV-2, RV-1 and RV-2, shown in separate panels. Significant differences in *gAI* between consecutive ROIs are marked with brackets (p < 0.05, bootstrap test; see Supp. Fig. S11 for single-subject results). Significant gradients in *gAI* are: *gAI_PT_* < *gAI_SMG_* < *gAI_PreG_* in LD-1, *gAI_HG_* < *gAI_msTS_* < *gAI_HG_* in LD-2, *gAI_PreG_* < *gAI_HG_* < *gAI_mSTG_* in LV-1, *gAI_HG_* < *gAI_mSTG_* < *gAI_aSTG_* in LV-2, and *gAI_mSTG_* < *gAI_aSTG_* in RV-2. In the left hemisphere, *gAI* gradually increases from early auditory regions to higher-order regions across the dorsal and ventral pathways. Similar patterns are also observed in the right hemisphere, although the gradients in *gAI* are less consistent across subjects.

#### Cortical representation of unattended speech

An important question regarding multi-speaker speech perception is to what extent unattended stimuli are represented in cortex. To address this question, here we investigated spectral, articulatory, and semantic representations of unattended stories during the cocktail-party task. We reasoned that if significant information about unattended speech is represented in a brain region, then features of unattended speech should explain significant variance in measured BOLD responses. To test this, we compared the prediction score of a combination model comprising the features of both attended and unattended stories (optimal convex combination) against the prediction score of an individual model comprising only the features of the attended story (see Methods). If the combination model significantly outperforms the individual model in an ROI, then the corresponding features of unattended speech are significantly represented in that ROI.

Figure 8 displays model performance when responses are predicted based on speech features from the attended story alone, and when they are instead predicted based on the optimally combined features from the attended and unattended stories. Results are shown for each ROI along the dorsal and ventral streams and in the left and right hemispheres (see Supp. Fig. S12a-e for single-subject results). Along the left (*HG*/*HS* → *PT* → *SMG* → (*POP*, *PreG*)) and right (*HG*/*HS* → *PT* → *SMG*) dorsal stream, spectral features of unattended speech are represented up to PT in LH and up to SMG in RH (p < 0.01), articulatory features are represented bilaterally up to PT (p < 0.05), whereas no semantic representation is apparent (p > 0.05). Along the left ventral stream (*HG*/*HS* → *mSTG* → *mSTS* → *MTG* and *HG*/*HS* → *mSTG* → *aSTG* → *PTR*), spectral and articulatory features are represented in HG/HS (p < 10^-4^), again with no semantic representation (p > 0.05). In the right ventral stream (*HG*/*HS* → *mSTG* → *mSTS* → *MTG* and *HG*/*HS* → *mSTG* → *aSTG* → *PTR*), spectral features are represented in HG/HS; articulatory features are represented up to mSTG (p < 0.05); and semantic features are represented only in mSTS (p < 0.05). These results indicate that cortical representations of unattended speech in multi-speaker environments extend from the spectral to the semantic level, albeit semantic representations are constrained to right parabelt auditory cortex (mSTS). Furthermore, representations of unattended speech are more broadly spread across the right hemisphere. Note that prior studies have reported response correlations and anatomical overlap between these belt/parabelt auditory regions and the reorienting attention system in the right-hemisphere (Corbetta et al. 2008; Vossel et al. 2014; Puschmann et al. 2017). Therefore, relatively broader representations of unattended speech in the right hemisphere might facilitate distractor detection and filtering during auditory attention tasks.

**Figure 8:**
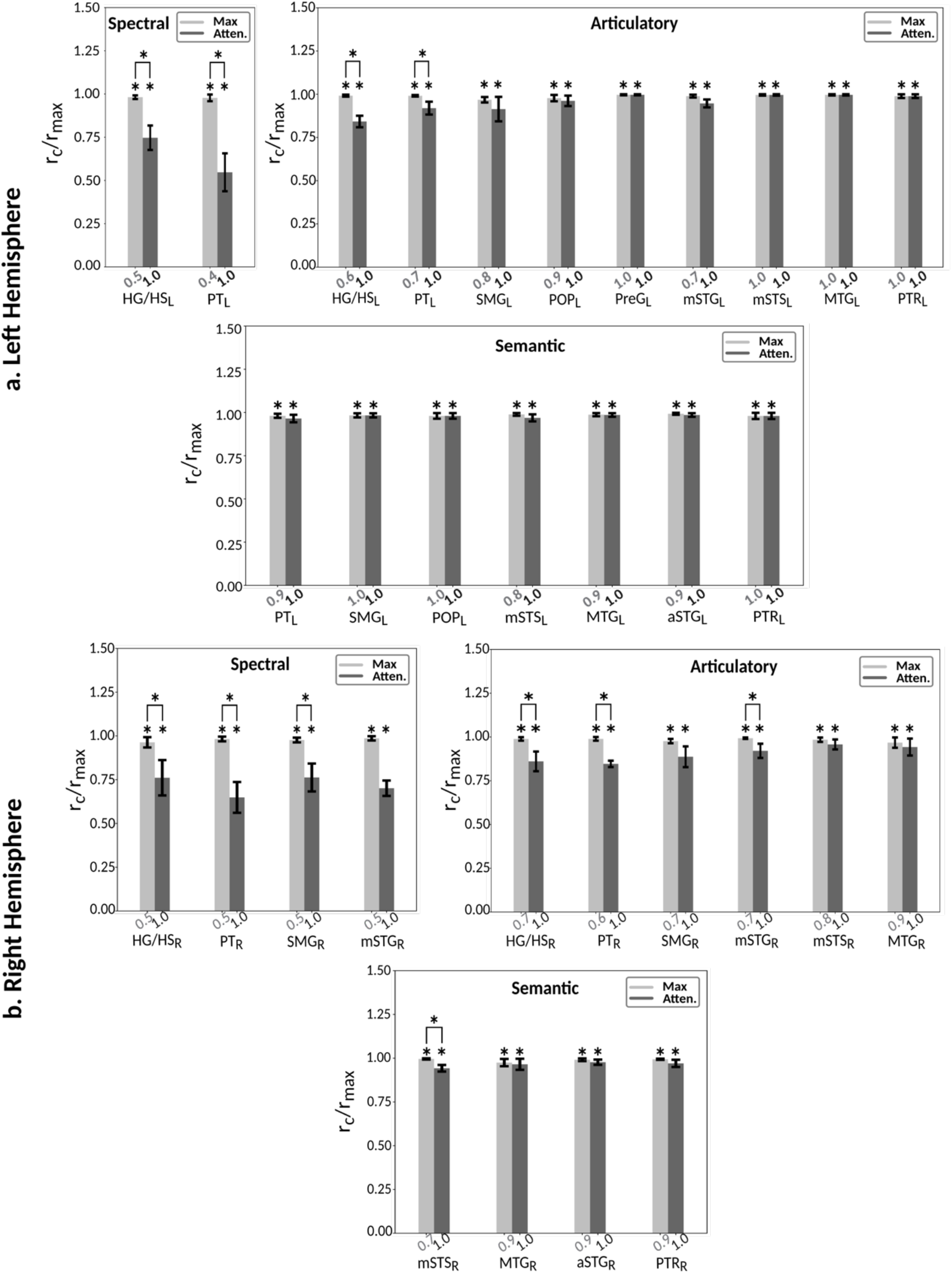
Representation of unattended speech. Passive-listening models were tested on cocktail-party data to assess representation of unattended speech during the cocktail-party task. Prediction scores were calculated separately for a combination model comprising features of both attended and unattended stories (*r_max_*: optimal convex combination) and an individual model only comprising features of the attended story (*r_a_*). Significant difference in prediction between the two models is an indication that BOLD responses carry significant information on unattended speech. Bar plots display normalized prediction scores (mean ± sem across subjects; combination model in light gray and individual model in gray). Significant scores are marked with * (p < 10^-4^, bootstrap test; see Supp. Fig. S12a-e for single-subject results), and significant differences are marked with brackets (p < 0.05). Prediction scores are displayed for ROIs in the dorsal and ventral streams, with significant selectivity for given model features. **a.** *Left hemisphere*. *Spectral representations* of unattended speech extend up to PT across the dorsal stream (*HG*/*HS* → *PT* → *SMG* → (*POP*, *PreG*)) and are constrained to HG/HS across the ventral stream (*HG*/*HS* → *mSTG* → *mSTS* → *MTG* and *HG*/*HS* → *mSTG* → *aSTG* → *PTR*). *Articulatory representations* of unattended speech extend up to PT across the dorsal stream and are constrained to HG/HS across the ventral stream. No *significant semantic representation* is apparent. **b.** *Right hemisphere*. *Spectral representations* of unattended speech extend up to SMG across the dorsal stream and are constrained to HG/HS across the ventral stream. *Articulatory representations* of unattended speech extend up to PT across the dorsal stream, and up to mSTG across the ventral stream. *Semantic representations* are found only in mSTS. These results suggest that processing of unattended speech is not constrained at spectral level but extends to articulatory and semantic level.

## Discussion

In this study, we investigated the effects of auditory attention on multi-level speech representations across cortex during a diotic cocktail-party task with naturalistic stimuli composed of spoken narratives. To assess baseline selectivity for multi-level speech features, we first fit spectral, articulatory, and semantic models using responses recorded during passive listening. We then quantified the complexity of intrinsic representations in each brain region. Next, we used fit models that reflect baseline selectivity for speech features to assess attentional modulation of speech representations. To do this, responses predicted using stimulus features of attended and unattended stories were compared with responses recorded during the cocktail-party task. This study is among the first to quantitatively characterize attentional modulations in multi-level speech representations of attended and unattended stimuli across speech-related cortex.

### Attentional Modulations

The effects of auditory attention on cortical responses have been primarily examined in the literature using controlled stimuli such as simple tones, melodies and isolated syllables or words (Alho et al. 1999; Jäncke et al. 2001, 2003; Lipschutz et al. 2002; Petkov et al. 2004; Johnson and Zatorre 2005; Degerman et al. 2006; Rinne et al. 2005, 2008, 2010; Woods et al. 2009, 2010; Paltoglou et al. 2009; Da costa et al. 2013; Seydell et al. 2014; Riecke et al. 2017). As such, less is known regarding how attention alters hierarchical representations of natural speech. Recent studies on this topic have mainly reported attentional modulations of low-level speech representations comprising speech-envelope and spectrogram features in early auditory and higher-order regions during the cocktail-party task (Ding and Simon 2012a, 2012b; Mesgarani and Chang 2012; Zion Golumbic et al. 2013; Puvvada and Simon 2017; Puschmann et al. 2019). Going beyond, here we have explored attentional modulations spanning from low-level spectral to high-level semantic features. While our results indicate that attentional modulations for articulatory and semantic representations distribute broadly across cortex, we find no consistent modulations for spectral representations in speech-related regions. Note that speech envelope and spectrogram features in natural speech carry intrinsic information about linguistic features including syllabic boundaries and articulatory features (Ding and Simon 2014; Liberto et al. 2015). These stimulus correlations can render it challenging to dissociate unique selectivity for articulatory versus spectral features. To minimize biases from potential stimulus correlations, here we leveraged a decorrelation procedure to obtain orthogonal spectral, articulatory, and semantic feature matrices for the stimulus. Therefore, the distinct modeling procedures for natural speech features might have contributed to the disparities between the current and previous studies on the existence of spectral modulations.

An important question regarding auditory attention is how the strength of attentional effects are distributed across cortex. A common view is that attentional modulations grow relatively stronger towards later stages of processing (Golumbic et al. 2013). Recent studies support this view by reporting bilaterally stronger modulations in frontal versus temporal cortex (Regev et al. 2019) and in non-primary versus primary auditory cortex (O’Sullivan et al. 2019). Adding to this body of evidence, we further show that attentional modulations gradually increase across the dorsal and ventral streams in the left hemisphere, as the complexity of speech representations grow. While a similar trend is observed across the right hemisphere, gradients in attentional modulation are less consistent in right belt and parabelt auditory regions including PT. Furthermore, attentional modulations are weaker in the right versus left hemisphere within these regions. Note that belt and parabelt regions are suggested to be connected to the right temporo-parietal junction (TPJ) during selective listening (Puschmann et al. 2017). TPJ is one of the central nodes in the reorienting attention system that monitors salient events to filter out distractors and help maintaining focused attention (Corbetta and Schulman 2002; Corbetta et al. 2008; Vossel et al. 2014). Hence less consistent gradients and relatively weaker attentional modulations in right belt and parabelt auditory regions might suggest a functional role for these regions in detecting salient events within the unattended stream during selective listening tasks.

Another central question regarding mechanisms of selective attention in a multi-speaker environment is how attentional modulations distribute across well-known dorsal and ventral streams. The dorsal stream that hosts articulatory representations is commonly considered to be involved in sound-to-articulation mapping (Hickok and Poeppel 2007, 2015; Rauschecker and Scott 2009; Friederici 2011; Rauschecker 2011). The motor-theory of speech perception suggests that dorsal articulatory representations carry information about articulatory gestures of the speaker to facilitate the listener’s comprehension (Liberman and Mattingly 1985; Hickok and Poeppel 2004; Davis and Johnsrude 2007; Scott et al. 2009b; Möttönen et al. 2013). Recent studies support this account by reporting enhanced activity in precentral gyrus and premotor cortex during challenging listening conditions (Osnes et al. 2011; Wild et al. 2012; Hervais-Adelman et al. 2012). In accordance with the motor theory of speech perception, here we find predominant articulatory selectivity and modulation due to selective listening in one end of the dorsal stream (PreG). These articulatory modulations might serve to increase sensitivity to the target speaker’s gestures to facilitate speech comprehension during difficult cocktail-party tasks (Wild et al. 2012).

In contrast to the dorsal stream, the ventral stream has been implicated in sound-to-meaning mapping (Hickok and Poeppel 2007, 2015; Rauschecker and Scott 2009; Friederici 2011; Rauschecker 2011). Compatible with this functional role, the ventral stream is suggested to transform acoustic representations of linguistic stimuli into object-based representations (Bizley and Cohen 2013). Here, we find that representational complexity of the speech features gradually increases across bilateral ventral stream, and semantic representations become dominant at the ends of it (bilateral PTR and MTG). In addition, speech level of attentional modulation also progresses across the ventral stream, and strong and predominant semantic modulations manifest towards later stages. Hence, the ventral stream might serve as a stage for interplay between bottom-up processing and top-down attentional modulation to gradually form auditory objects during selective listening (Bizley and Cohen 2013; Shinn-Cunningham et al. 2017; Rutten et al. 2019).

### Representation of the unattended speech

Whether unattended speech is represented in cortex during selective listening and if so, at what feature levels its representations are maintained are crucial aspects of auditory attention. Behavioral accounts suggest that unattended speech is primarily represented at the acoustic level (Cherry 1953; Broadbent 1958). Corroborating these accounts, recent electrophysiology studies have identified acoustic representations of unattended speech localized to auditory cortex (Ding and Simon 2012a, 2012b; Zion Golumbic et al. 2013; Puvvada and Simon et al. 2017; Brodbeck et al. 2018b; O’Sullivan et al. 2019; Puschman et al. 2019). In contrast, here we find that acoustic representations of unattended speech extend beyond the auditory cortex as far as SMG in the right dorsal stream. Because SMG partly overlaps with the reorienting attention system, unattended speech representations in this region might contribute to filtering of distractors during the cocktail-party task (Corbetta et al. 2008; Vossel et al. 2014).

A more controversial issue is whether unattended speech representations carry information at the linguistic level (Driver 2001; Lavie 2005; Boulenger et al. 2010; Bronkhorst 2015; Kidd and Colburn 2017). Prior studies on this issue are split between those suggesting the presence (Wild et al. 2012; Evans et al. 2016) versus absence (Sabri et al. 2008; Brodbeck et al. 2018b) of linguistic representations. Here, we find that articulatory representations of unattended speech extend up to belt/parabelt auditory areas in the bilateral dorsal stream and the right ventral stream. We further find semantic representation of unattended speech in the right ventral stream (mSTS). These linguistic representations of unattended speech are naturally weaker than those of attended speech, and they are localized to early-to-intermediate stages of auditory processing. Our findings suggest that unattended speech is represented at the linguistic level prior to entering the broad semantic system where full selection of the attended stream occurs (Bregman 1994; Pulvermüller and Shtyrov 2006; Relander et al. 2009; Näätänen et al. 2011; Rämä et al. 2012; Bronkhorst 2015; Ding et al. 2018). Overall, these linguistic representations might serve to direct exogenous triggering of attention to salient features in unattended speech (Moray 1959; Treisman 1960, 1964; Wood and Cowan 1995; Driver 2001; Bronkhorst 2015). Meanwhile, attenuated semantic representations in the ventral stream might facilitate semantic priming of the attended stream by relevant information in the unattended stream (Lewis 1970; Driver 2001; Rivenez et al. 2006).

### Future work

Reliability of statistical assessments in neuroimaging depends on two main factors: sample size and amount of data collected per subject. Given experimental constraints, it is difficult to increase both factors in a single study. In this unavoidable trade-off, a common practice in fMRI studies is to collect a relatively limited dataset from more subjects. This practice prioritizes across-subject variability over within-subject variability, at the expense of individual-subject results. Diverting away from this practice, we collected a larger amount of data per subject to give greater focus to reliability in single subjects. This choice is motivated by the central aims of the voxel-wise modeling (VM) approach. The VM framework aims to sensitively measure tuning profiles of single voxels in individual subjects. For the natural speech perception experiments conducted here, the tuning profiles were characterized over three separate high-dimensional spaces containing hundreds of acoustic and linguistic features. To maximize sensitivity of VM models, we conducted extensive experiments in each individual subject to increase the amount and diversity of fMRI data collected. This design enhanced the quality of resulting VM models and reliability of individual-subject results. Indeed, here we find highly uniform results across individual subjects, suggesting that the reported effects are highly robust. That said, across subject and across language variability might occur in diverse, multi-lingual cohorts. Assessment of attentional effects on speech representations in these broader populations remains important future work.

In the current study, subjects were presented continuous natural speech stimuli. In the passive-listening task, they were instructed to vigilantly listen to the presented story. Our analyses reveal that BOLD responses in large swaths of language-related areas can be significantly predicted by voxel-wise models comprising spectral, articulatory and semantic speech features. Moreover, the spectral, articulatory and semantic representations mapped in single subjects are highly consistent across subjects. Therefore, these results suggest that the participants performed reasonably well in active listening of single-speaker stories. In the cocktail-party experiment, subjects were instead instructed to attentively listen to one of two speakers. Our analyses in this case reveal broad attentional modulations in representation of semantic information across cortex, in favor of the target speaker. Semantic features of natural speech show gradual variation across time compared to low-level spectral information. Therefore, this finding suggests that subjects also performed well during the sustained attention tasks in the cocktail-party task. That said, we cannot rule out momentary shifts in attention away from the target speaker. If momentary shifts towards the unattended speaker are frequent, they might increase the proportion of unattended speech information that BOLD responses carry. In turn, this might have contributed to the strength of unattended speech representations that we measured during the cocktail-party task. Post-scan questionnaires that assess participants’ comprehension of attended and unattended stories are a common control for task execution (Regev et al. 2018). However, post-scan memory controls cannot guarantee lack of momentary attention shifts that typically last less than 200 ms (Spence and Driver 1997). On the other hand, implementing frequent controls during the scan itself would disrupt the naturalistic experiment flow and efficiency. It is therefore challenging to experimentally monitor momentary attentional shifts (Bronkhorst, 2015). To assess the influence of momentary shifts during sustained-attention tasks, future studies are warranted leveraging more controlled speech stimuli with systematic variations in the salience and task relevance of non-target stimuli (Corbetta et al. 2008; Parmentier et al. 2014; Bronkhorst 2015).

In the current study, we find that attention strongly alters semantic representations in favor of the target stream across frontal and parietal cortices. This is in alignment with previous fMRI studies that found attentional response modulations in frontal and parietal regions (Ikeda et al. 2010; Hill and Miller 2010; Regev et al. 2019). That said, an important question is whether these modulations predominantly reflect enhanced bottom-up processing of attended speech or top-down attentional control signals (Corbetta and Shulman 2002; Seydell-Greenwald et al. 2014). Note that we find broad semantic representations across frontal and parietal cortices during the passive-listening experiment, in the absence of any demanding attentional tasks. Furthermore, a recent study suggests that various semantic categories are differentially represented in these higher-level cortical regions (Huth et al. 2016). Taken together, these findings imply that semantic modulations in fronto-parietal regions can be partly attributed to bottom-up effects. Yet, it is challenging to disentangle bottom-up and top-down contributions in fMRI studies due to the inherently limited temporal resolution. Future studies are warranted to shed light on this issue by combining the spatial sampling capability of fMRI with high temporal resolution of electrophysiology methods (Cukur et al. 2013; de Heer et al. 2017).

## Conclusion

In sum, our results indicate that attention during a diotic cocktail-party task with naturalistic stimuli gradually selects attended over unattended speech across both dorsal and ventral processing pathways. This selection is mediated by representational modulations for linguistic features. Despite broad attentional modulations in favor of the attended stream, we still find that unattended speech is represented up to linguistic level in the regions that overlap with the reorienting attention system. These linguistic representations of unattended speech might facilitate attentional reorienting and filtering during natural speech perception. Overall, our findings provide comprehensive insights on attentional mechanisms that underlie the ability to selectively listen to a desired speaker in noisy multi-speaker environments.

## Notes

The authors thank Jack L. Gallant, Wendy de Heer and Ümit Keleş for assistance in various aspects of this research. *Conflict of interest*: None.

